# Novel cerebello-amygdala connections provide missing link between cerebellum and limbic system

**DOI:** 10.1101/2022.02.07.479043

**Authors:** Se Jung Jung, Ksenia Vlasov, Alexa D’Ambra, Abhijna Parigi, Mihir Baya, Edbertt Paul Frez, Jacqueline Villalobos, Marina Fernandez-Frentzel, Maribel Anguiano, Yoichiro Ideguchi, Evan G. Antzoulatos, Diasynou Fioravante

## Abstract

The cerebellum is emerging as a powerful regulator of cognitive and affective processing and memory in both humans and animals and has been implicated in affective disorders. How the cerebellum supports affective function remains poorly understood. The short-latency (just a few ms) functional connections that were identified between the cerebellum and amygdala -a structure crucial for the processing of emotion and valence-more than 4 decades ago raise the exciting, yet untested, possibility that a cerebellum-amygdala pathway communicates information important for emotion. The major hurdle in rigorously testing this possibility is the lack of knowledge about the anatomy and functional connectivity of this pathway. Our initial anatomical tracing studies in mice excluded the existence of a direct monosynaptic pathway between cerebellum and amygdala. Using transneuronal tracing techniques, we have identified a novel disynaptic pathway that connects the cerebellar output nuclei to the basolateral amygdala. This pathway recruits the understudied intralaminar thalamus as a node. Using ex vivo optophysiology and super-resolution microscopy, we provide the first evidence for the functionality of the pathway, thus offering a missing mechanistic link between the cerebellum and amygdala. This discovery provides a connectivity blueprint between the cerebellum and a key structure of the limbic system. As such, it is the requisite first step toward obtaining new knowledge about cerebellar function in emotion, thus fundamentally advancing understanding of the neurobiology of emotion, which is perturbed in mental and autism spectrum disorders.

## 1 Introduction

The cerebellum is increasingly recognized as a regulator of limbic functions ^1–8^. The human cerebellum is activated in response to aversive or threatening cues, upon remembering emotionally charged events, and during social behavior, reward-based decision making and violation of expectation ^9–18^. Consistent with this, deficits in cerebellar function are associated with impaired emotional attention and perception, as seen in depression, anxiety, schizophrenia and post-traumatic stress disorder ^19–22^, as well as cognitive and emotional disturbances collectively known as cerebellar cognitive affective syndrome ^23^. Animal models have recapitulated some of these findings, with selective mutations, damage or inactivation of the rodent cerebellum resulting in altered acquisition or extinction of learned defensive responses, and impaired social and goal-directed behavior, without motor deficits ^24–33^.

The limited understanding of the anatomical and functional circuits that connect the cerebellum to limbic centers has impeded mechanistic insight into the neural underpinnings of cerebellar limbic functions, which have begun to be dissected only recently ^30–32,34^. Moreover, a neuroanatomical substrate for the functional connections between the cerebellum and a key affective center, the amygdala ^35^, has yet to be provided, even though these connections were observed more than 40 years ago ^36–38^. The purpose of the present work was to generate a mesoscale map of functional neuroanatomical connectivity between the cerebellum and amygdala. We focused on connections between the deep cerebellar nuclei (DCN), which give rise to most cerebellar output pathways ^39^, and the basolateral amygdala (BLA), which is known to process affect-relevant salience and valence information ^35,40,41^, and which was targeted in the early electrophysiological studies of Heath et al. ^37,38^.

## 2 Methods

### 2.1 Mice

C57Bl/6J mice of both sexes were used in accordance with National Institute of Health guidelines. All procedures were reviewed and approved by the Institutional Animal Care and Use Committee of the University of California, Davis. Mice were maintained on a 12-hr light/dark cycle with ad libitum access to food and water. For anatomical tracing experiments, postnatal day P45-65 (at the time of injection) mice were used (N = 13 mice). For slice optophysiology, P18-25 (at the time of injection) mice were used. (Fig. 3: N = 14 mice; Fig. 5: N = 5 mice; Fig. 6: N = 7 mice).

### 2.2 Virus and tracer injections

For stereotaxic surgeries, mice were induced to a surgical plane of anesthesia with 5% isoflurane and maintained at 1-2% isoflurane. Mice were placed in a stereotaxic frame (David Kopf Instruments, Tujunga, California) on a feedback-controlled heating pad. Following skin incision, small craniotomies were made above the target regions with a dental drill. The following coordinates (in mm) were used (from bregma): for medial DCN - 6.4 AP, ± 0.75 ML, −2.2 DV; for interposed DCN: −6.3 AP, ± 1.6 ML, −2.2 DV; for lateral DCN: −5.7 AP, ± 2.35 ML, −2.18 DV. For basolateral amygdala: −0.85 AP, ± 3.08 ML, −4.5 DV. For limbic thalamus: −0.85 AP, ± 0.3 ML, −3.3 DV, and −1.2 AP, ± 0.5 ML, −3.5 DV. A small amount of tracer (50 - 100 nl for DCN, 300 - 500 nl for thalamus) was pressure-injected in the targeted site with a UMP3-1 ultramicropump (WPI, Sarasota, FL) and glass pipettes (Wiretrol II, Drummond) (tip diameter: 25-50 μm) at a rate of 30 nl/min. The pipette was retracted 10 min after injection, the skin was sutured (Ethilon P-6 sutures, Ethicon, Raritan, NJ) and/or glued (Gluture, Abbott labs, Abbott Park, IL) and animal was allowed to recover completely prior to returning to the home cage. Preoperative analgesia consisted of a single administration of local lidocaine (VetOne, MWI, Boise, ID; 1 mg/kg) and Meloxicam (Covetrus, Portland, ME; 5 mg/kg), both SC. Postoperative analgesia consisted of a single administration of Buprenex (AmerisourceBergen Drug Corp, Sacramento, CA; 0.1 mg/kg) and Meloxicam 5 mg/kg, both SC, followed by Meloxicam at 24 and 48 hr. The following adeno-associated viruses (AAV) and tracers were used: AAV8-CMV-TurboRFP (UPenn Vector Core, 1.19*10^14 gc/ml), AAV9-CAG-GFP (UNC Vector Core, 2×10^12 gc/ml), AAV2-retro-CAG-GFP (Addgene, 7×10^12 gc/ml), AAV2-retro-AAV-CAG-tdTomato (Addgene, 7×10^12 gc/ml), Cholera toxin subunit B CF-640 (Biotium, 2 mg/ml, 100 nl), AAV1-hSyn-Cre-WPRE-hGH (Addgene, 10^13 gc/ml, diluted 1:5), AAV5-CAG-FLEX-tdtomato (UNC Viral Core, 7.8*10^12 gc/ml, diluted 1:5), AAV9-EF1a-DIO-hChR2(H134R)-EYFP (Addgene, 1.8*10^13 gc/ml, diluted 1:10), AAV2-hSyn-hChR2(H134R)-EYFP (UNC Vector Core, 5.6×10^12 gc/ml, diluted 1:2). Three-five weeks were allowed for viral expression/labelling.

### 2.3 Histology and imaging

Following deep anesthesia (anesthetic cocktail: 100 mg/kg ketamine, 10 mg/kg xylazine, 1 mg/kg acepromazine, IP) mice were paraformaldehyde-fixed (4% paraformaldehyde in 0.1 M phosphate buffer, pH 7.4, EMS Diasum, Hatfield, PA) through transcardial perfusion. Brains were post-fixed overnight, cryo-protected with 30% sucrose in PBS and sliced coronally on a sliding microtome at 60-100 μm thickness. Slices were mounted on slides with Mowiol-based mounting media and scanned using an Olympus VS120 Slide Scanner (Olympus, Germany) (resolution with 10x/0.4 N.A. lens at 488 nm: 645 nm in x,y). For immunohistochemistry, slices were blocked with 10% normal goat serum (NGS, Millipore, Burlington, MA) in PBST (0.3% Triton X-100 in PBS) for 1 h. Slices were incubated with primary antibodies (anti-Cre IgG1, Millipore, 1:1000; anti-NEUN, Cell Signaling, Danvers, MA, 1:1000; anti-vGLUT2, Synaptic Systems, Goettingen, Germany, 1:700; anti-PSD-95, Neuromab, Davis, CA, 1:500) in 2% NGS-PBST overnight at 4°C. After 4 x 20-min rinses with PBST, secondary antibodies (Alexa fluor-568 goat anti-mouse 1:1000 IgG1; Alexa fluor-488 goat anti-rabbit 1:1000; Dylight-405 goat anti-guinea pig 1:200; Alexa fluor-647 goat anti-mouse 1:1000 IgG2a; Life Technologies, Carlsbad, CA) were applied in 2% NGS-PBST for 1-2 h at room temperature. Following another round of rinses, slices were mounted on slides with Mowiol and scanned on an LSM800 confocal microscope with Airyscan (resolution with 63x/1.4 N.A. oil lens at 488 nm: 120 nm in x,y, 350 nm in z) (Zeiss, Germany). Maximal projections of optical z-stacks were obtained with Zen software (Zeiss) and used for analysis.

### 2.4 Preparation of brain slices for electrophysiology

Mice of either sex were anesthetized through intraperitoneal injection of ketamine/xylazine/acepromazine anesthetic cocktail and transcardially perfused with ice-cold artificial cerebrospinal fluid (aCSF; in mM: 127 NaCl, 2.5 KCl, 1.25 NaH_2_PO_4_, 25 NaHCO_3_, 1 MgCl_2_, 2 CaCl_2_, 25 glucose; supplemented with 0.4 sodium ascorbate and 2 sodium pyruvate; ~310 mOsm). Brains were rapidly removed, blocked, and placed in choline slurry (110 choline chloride, 25 NaHCO_3_, 25 glucose, 2.5 KCl, 1.25 NaH_2_PO_4_, 7 MgCl_2_, 0.5 CaCl_2_, 11.6 sodium ascorbate, 3.1 sodium pyruvate; ~310 mOsm). Coronal sections (250 μm) containing the thalamus were cut on a vibratome (Leica VT1200S) and allowed to recover in aCSF at 32°C for 25 min before moving to room temperature until further use. All solutions were bubbled with 95% O_2_-5% CO2 continuously. Chemicals were from Sigma.

### 2.5 Electrophysiology

Slices were mounted onto poly-l-lysine-coated glass coverslips and placed in a submersion recording chamber perfused with aCSF (2-3 ml/min) at near physiological temperature (30-32°C). Whole-cell voltage-clamp recordings were made from tdTomato+ (Figs. 3,5) or CtB+ (Fig. 6) cells in the thalamus using borosilicate glass pipettes (3-5 MΩ) filled with internal solution containing (in mM): CsMSO_3_ 120, CsCl 15, NaCl 8, TEA-Cl 10, HEPES 10, EGTA 0.5, QX314 2, MgATP 4 and NaGTP 0.3, biocytin 0.3. Recordings were acquired in pClamp11 using a Multiclamp 700B amplifier (Molecular Devices, San Jose, CA), digitized at 20 kHz and low-pass filtered at 8 kHz. Membrane potential was maintained at −70 mV. Series resistance and leak current were monitored and recordings were terminated if either of these parameters changed significantly. Optical stimulation of ChR2+ fibers surrounding tdTomato+ or CtB+ thalamic neurons was performed under a 60x water immersion lens (1.0 N.A.) of an Olympus BX51W microscope, using an LED system (Excelitas X-cite; or Prizmatix UHP-T) mounted on the microscope and driven by a Master9 stimulator (AMPI). Optical stimulation consisted of 488 nm light pulses (1-5 ms duration). Power density was set to 1.5-2x threshold (max: 0.25 mW/mm^2^). A minimum of 5 response-evoking trials (inter-trial interval: 60 s) were delivered and traces were averaged. To confirm monosynaptic inputs, action potentials were blocked with TTX (1 μM), followed by TTX+ 4AP (100 μM) to prolong ChR2-evoked depolarization ^42^.

### 2.6 Data analysis

Analysis of ex vivo recordings was performed using custom MATLAB R2019b scripts (MathWorks, Natick, MA). Postsynaptic current (PSC) amplitude was computed from the maximum negative deflection from baseline within a time window (2.5 - 40 ms) from stimulus onset. Onset latency was measured at 10% of peak amplitude. Cell location was confirmed through biocytin-streptavidin Alexa fluor staining. For slice registration the Paxinos Brain Atlas (Paxinos and Franklin, 2001) and the Allen Brain Atlas (ABA_v3) were used. Location of injection sites was identified and experiments were excluded if there was spill into neighboring nuclei. Cell counting and immunofluorescence intensity analyses were done by raters naïve to the experimental hypotheses using ImageJ (Fiji, National Institutes of Health, Bethesda, Maryland) and Abode Illustrator. Statistical analysis was performed in Matlab (Mathworks) and Prism (GraphPad), with significance set at p < 0.05.

## 3 Results

### 3.1 Putative disynaptic pathways between cerebellar nuclei and BLA through the limbic thalamus

Given that microstimulation of DCN elicits short-latency responses in the BLA ^36–38^, we hypothesized that an anatomical pathway exists between the two regions that involves at most 2 synapses. Initial anatomical tracing experiments did not support a direct DCN-BLA connection (not shown). We therefore performed simultaneous injections of an anterograde tracer virus (AAV8-CMV-TurboRFP) in the DCN and a retrograde tracer virus (AAV2-retro-CAG-GFP) in the BLA (**Fig. 1A,B**) to identify potential regions of overlap. In epifluorescence images of brain slices across different animals (N = 8), the limbic thalamus consistently emerged as a prominent site of overlap (**Fig. 1C**). We use the term “limbic thalamus” to refer to a collection of non-sensorimotor thalamic nuclei, including the mediodorsal (MD), midline and intralaminar (IL) nuclei, with diverse projections to cortical (mainly medial prefrontal) and/or subcortical limbic structures ^43–46^. Registration of images to the Allen Brain Atlas localized BLA-projecting thalamic neurons in multiple nuclei of the limbic thalamus (**Fig. 1D**), in agreement with known connectivity patterns ^46–49^. Visual inspection of diffraction-limited epifluorescence images identified overlapping DCN axonal projections and BLA-projecting neurons in several (but not all) of these thalamic nuclei, including the parafascicular (PF) n. and subparafascicular area (SPA), the centromedial (CM) and MD nuclei, and other midline nuclei (**Fig. 1E**). Injection of the tracer cholera toxin subunit B (CtB)-CF640 in the limbic thalamus retrogradely labeled neurons in DCN (**Fig. 1F**), confirming the DCN-limbic thalamus connectivity.

**Figure 1.**
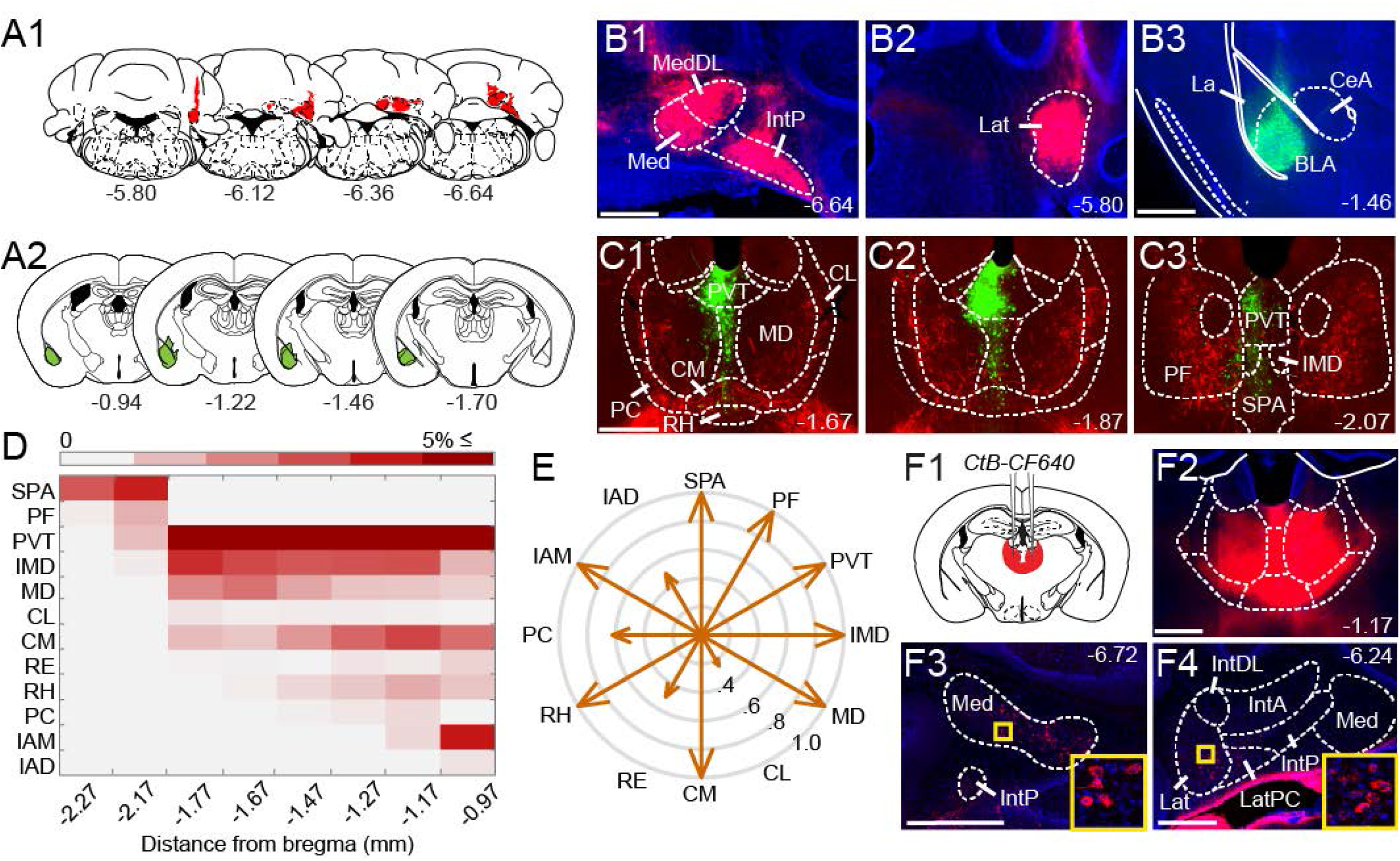
Anatomical tracing uncovers putative disynaptic pathways from cerebellum to basolateral amygdala. **A,** Unilateral injection sites for anterograde viral tracer in DCN (**A1**, *red*) and retrograde viral tracer in BLA (**A2**, *green*). **B,** Mosaic epifluorescence images of injection sites in DCN (**B1-2**) and BLA (**B3**). **C,** Mosaic epifluorescence images of overlapping DCN axons (red) and BLA-projecting neurons (green) in limbic thalamus. **D,** Relative distribution of BLA-projecting neurons across nuclei of the limbic thalamus, normalized to the total number of labeled neurons and averaged across experiments, as a function of distance from bregma. Antero-posterior coordinates for each nucleus are given in **Table 1**. **E,** Quantification of overlap between DCN axons and BLA-projecting thalamic neurons. Arrow length in radar plot indicates proportion (0-1) of experiments with overlap in each thalamic nucleus. **F1,F2,** Schematic and confocal image of injection site for retrograde tracer CtB CF-640 in limbic thalamus. **F3,F4,** CtB-labeled projection neurons (red) in DCN at different distances from bregma. Insets show high-magnification images of areas in yellow squares. For all panels, numbers denote distance (in mm) from bregma. Scale bars: 500 μm.

### 3.2 Transneuronal anatomical tracing and optophysiology establish synaptic connectivity between cerebellar nuclei and limbic thalamus

To spatially resolve synaptic connectivity between DCN and BLA-projecting thalamic nuclei, we adopted an AAV-based transneuronal approach ^50^. AAV1-Cre in presynaptic neurons is known to propagate across the synapse and induce expression of a floxed tag in postsynaptic neurons, thus identifying synaptic partners (**Fig. 2A**). We injected AAV1-Cre bilaterally in DCN and AAV-FLEX-tdTomato in thalamus and quantified the relative distribution of tdTomato+ neurons in intralaminar and midline thalamic nuclei. Injection coverage for DCN was indicated by Cre immunofluorescence (**Fig. 2B1,2**) and included all cerebellar nuclei. Great care was taken to avoid spill to extracerebellar areas, which resulted in denser coverage of caudal DCN (**Fig. 2B3**). TdTomato+ neurons were observed throughout the limbic thalamus, confirming adequate coverage, and extended into ventromedial nuclei (**Fig. 2C**), which served as positive control ^51,52^. Averaging the relative distribution of tdTomato+ neurons across five successful experiments revealed that the intralaminar cluster, comprised of centrolateral (CL), paracentral (PC), CM, and PF nuclei ^47^, and MD nucleus encompassed most (~95%) tagged neurons (**Fig. 2C3**), suggesting that these nuclei reliably receive most cerebellar inputs to limbic thalamus.

**Table 1.**
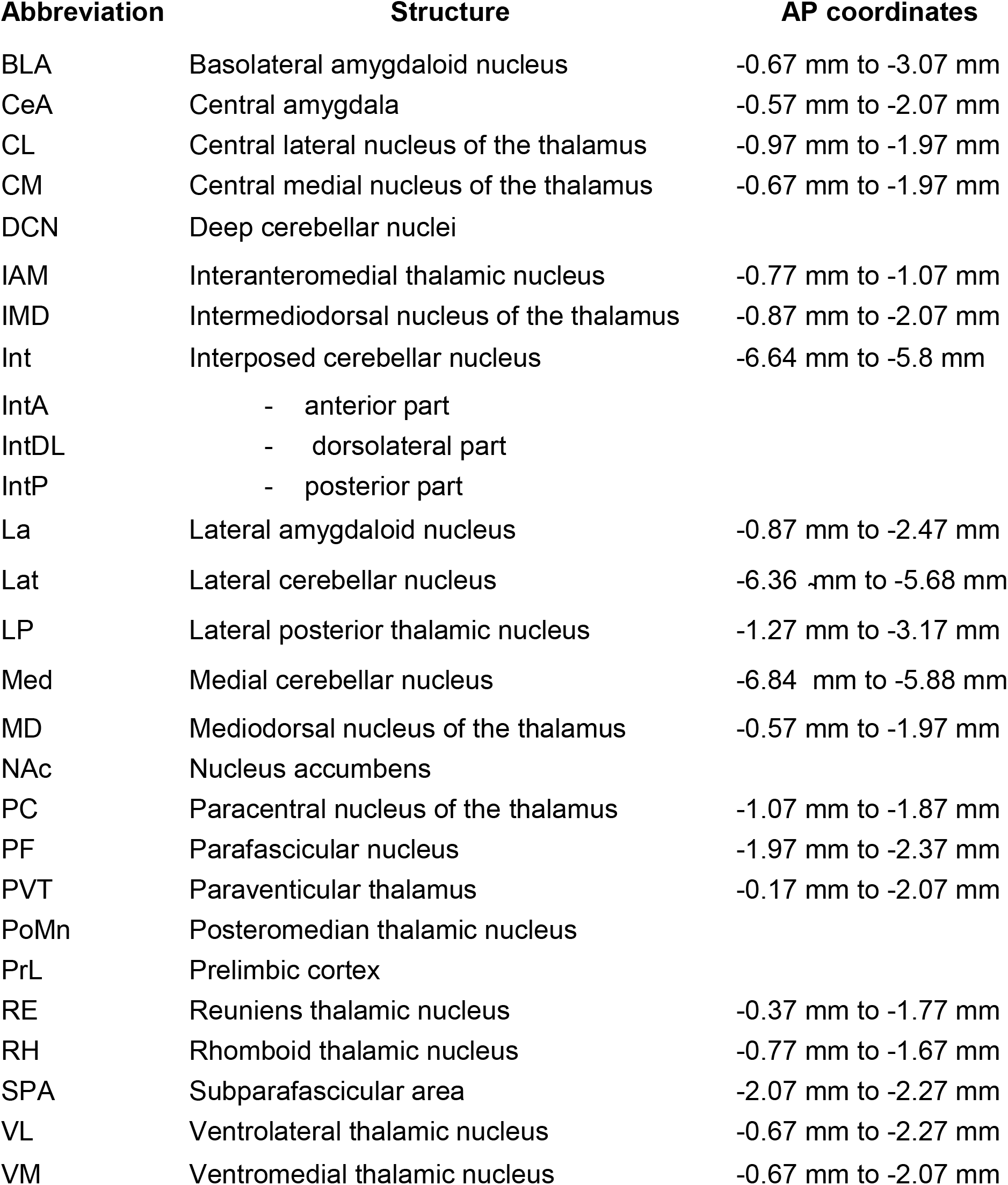
Anatomical abbreviations (in alphabetical order) and antero-posterior coordinates (in mm, from bregma)

| Abbreviation | Structure | AP coordinates |
| --- | --- | --- |
| BLA | Basolateral amygdaloid nucleus | -0.67 mm to -3.07 mm |
| CeA | Central amygdala | -0.57 mm to -2.07 mm |
| CL | Central lateral nucleus of the thalamus | -0.97 mm to -1.97 mm |
| CM | Central medial nucleus of the thalamus | -0.67 mm to -1.97 mm |
| DCN | Deep cerebellar nuclei |  |
| IAM | Interanteromedial thalamic nucleus | -0.77 mm to -1.07 mm |
| IMD | Intermediodorsal nucleus of the thalamus | -0.87 mm to -2.07 mm |
| Int | Interposed cerebellar nucleus | -6.64 mm to -5.8 mm |
| IntA | - anterior part |  |
| IntDL | - dorsolateral part |  |
| IntP | - posterior part |  |
| La | Lateral amygdaloid nucleus | -0.87 mm to -2.47 mm |
| Lat | Lateral cerebellar nucleus | -6.36 mm to -5.68 mm |
| LP | Lateral posterior thalamic nucleus | -1.27 mm to -3.17 mm |
| Med | Medial cerebellar nucleus | -6.84 mm to -5.88 mm |
| MD | Mediodorsal nucleus of the thalamus | -0.57 mm to -1.97 mm |
| NAc | Nucleus accumbens |  |
| PC | Paracentral nucleus of the thalamus | -1.07 mm to -1.87 mm |
| PF | Parafascicular nucleus | -1.97 mm to -2.37 mm |
| PVT | Paraventricular thalamus | -0.17 mm to -2.07 mm |
| PoMn | Posteromedian thalamic nucleus |  |
| PrL | Prelimbic cortex |  |
| RE | Reuniens thalamic nucleus | -0.37 mm to -1.77 mm |
| RH | Rhomboid thalamic nucleus | -0.77 mm to -1.67 mm |
| SPA | Subparafascicular area | -2.07 mm to -2.27 mm |
| VL | Ventrolateral thalamic nucleus | -0.67 mm to -2.27 mm |
| VM | Ventromedial thalamic nucleus | -0.67 mm to -2.07 mm |

**Figure 2.**
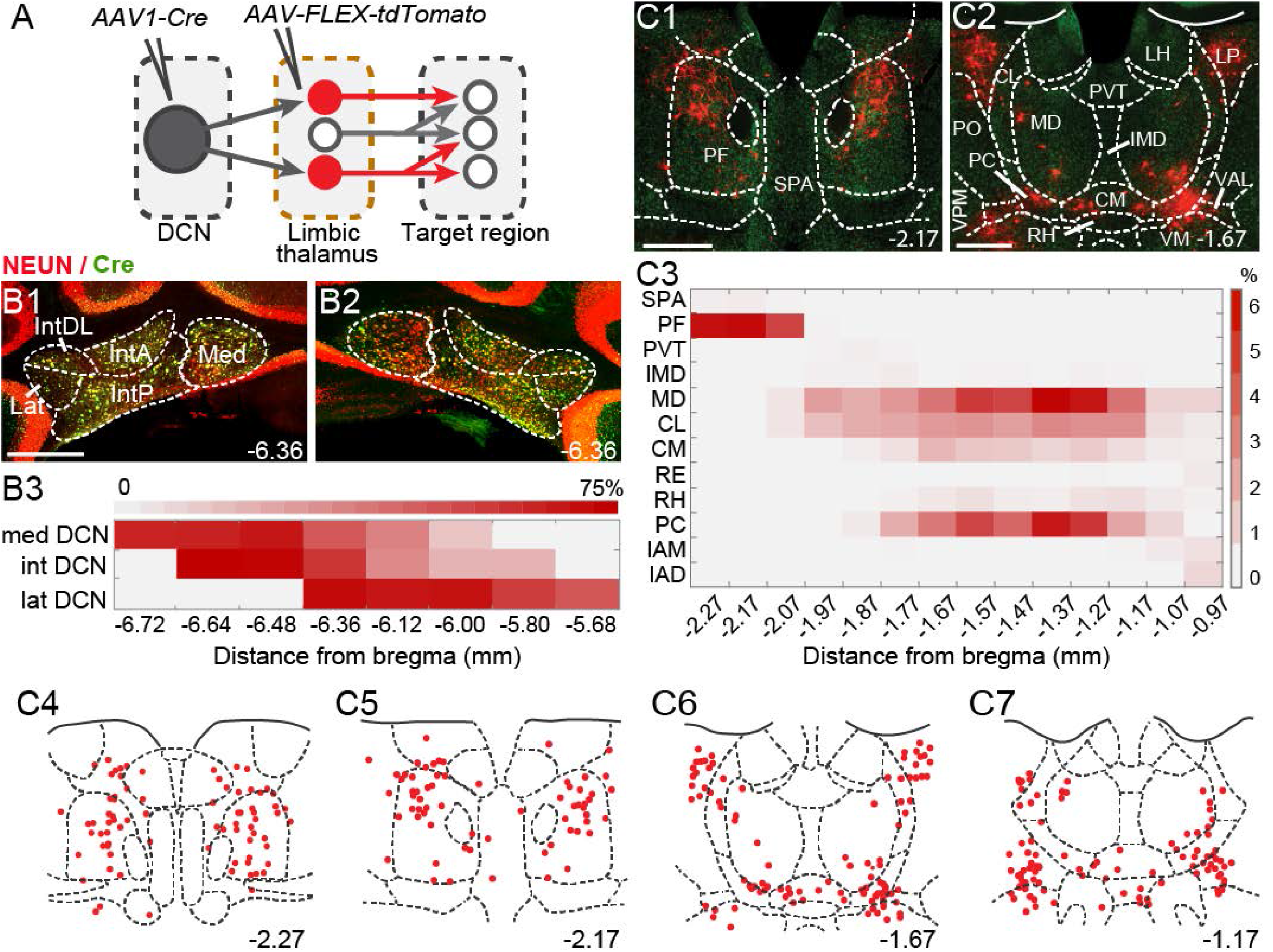
The intralaminar and mediodorsal nuclei are major cerebellar postsynaptic targets in limbic thalamus. **A,** Schematic of experimental approach for disynaptic pathway tracing. **B1-B2,** Example images of bilateral Cre expression in DCN. Red: immunofluorescence for NeuN neural marker; Green: anti-Cre immunoreactivity; Yellow: merge. **B3,** Heatmap of Cre immunofluorescence in DCN, normalized to NeuN signal and averaged across experiments, as a function of distance (in mm) from bregma. **C1,C2,** Example images of thalamic neurons conditionally expressing tdTomato (red) upon transneuronal transfer of Cre from cerebellar presynaptic axons. Green: NeuN immunofluorescence. **C3,** Heatmap of relative distribution of tdTomato+ neurons across thalamic nuclei, normalized to total number of labeled neurons and averaged across experiments, as a function of distance from bregma. **C4-C7,** Example registration of tdTomato+ neurons to the Allen mouse brain atlas. Numbers at bottom denote distance (in mm) from bregma. Antero-posterior coordinates for each nucleus can be found in **Table 1**. Scale bars: 500 μm.

To confirm that the thalamic targets identified with the transneuronal Cre method receive cerebellar synaptic input, we performed optophysiological experiments in acute thalamic slices from mice injected with AAV1-Cre in the DCN and AAV-FLEX-tdTomato in the thalamus (**Fig. 3A**). To activate cerebellar inputs, channelrhodopsin (ChR2-H134R) was conditionally expressed in DCN through AAV-DIO-ChR2-EYFP injection. DCN axonal projections were stimulated in the thalamus with 488-nm light pulses applied through the objective. Light-evoked synaptic responses were monitored in whole-cell voltage-clamp recordings (V_m_ = −70 mV) from thalamic neurons, which were selected based on tdTomato expression, their anatomical location and position in the slice, i.e., surrounded by ChR2-EYFP-expressing axons. In all thalamic nuclei examined (n = 29 cells), light stimulation elicited synaptic responses (mean response in pA: IL: 311.7 ± 100; MD: 105.7 ± 32.3; midline: 565.8 ± 209.8; VM/VPM: 347.5 ± 112.3; LP: 91.8 ± 2.7) (**Fig. 3B1**) with short latencies (mean latency in ms: IL: 2.5 ± 0.28; MD: 3.3 ± 0.6; midline: 4.2 ± 0.7; VM/VPM: 3.2 ± 0.2; LP: 2.9 ± 0.8) (**Fig. 3B2**). These data support the specificity of the anatomical connectivity and establish the existence of active DCN terminals (as opposed to just passing axons) across limbic thalamus.

**Figure 3.**
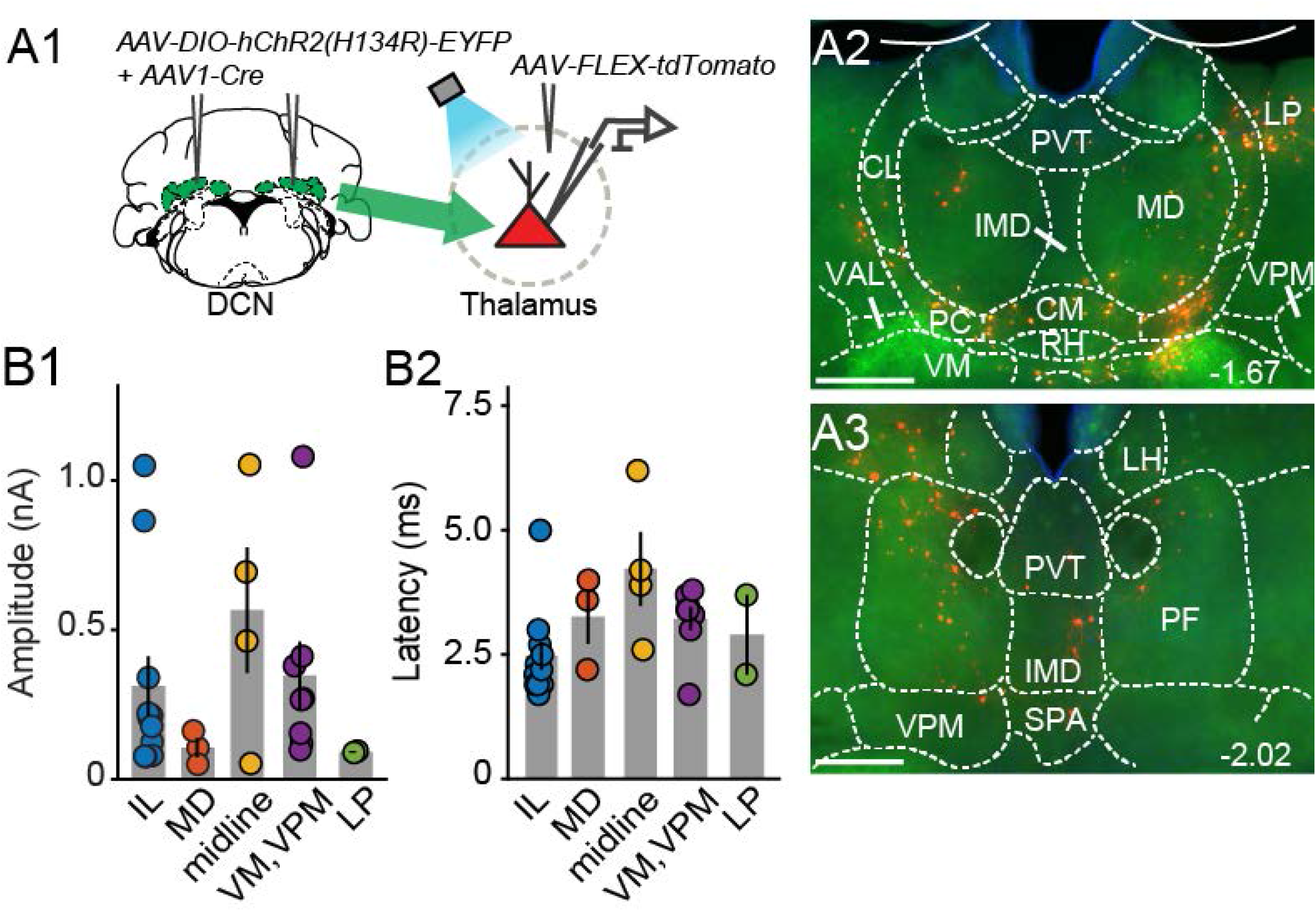
Electrophysiological validation of virally-identified cerebello-thalamic connectivity. **A1,** Schematic of experimental approach for ex vivo optophysiology. **A2,A3,** Epifluorescence images of anterior (A2) and posterior (A3) thalamic slices acutely prepared for recordings. DCN input-receiving neurons are tdTomato+. Scale bars: 500 μm. **B,** Average (± SEM) amplitude (**B1**) and onset latency (**B2**) of ChR2-evoked synaptic currents as a function of recording location in the thalamus. Intralaminar (IL) group: CL, PC, CM and PF; midline group: IMD and RH.

### 3.3 Thalamic neurons receiving cerebellar input project to BLA

If the thalamus is a functional node of the disynaptic DCN-BLA circuit, then we would expect to find axons of DCN input-receiving thalamic neurons in BLA. To this end, we imaged BLA-containing slices from transsynaptic Cre experiments (**Fig. 4A**). We detected tdTomato+ axons at several antero-posterior distances from bregma (**Fig. 4B1–B6**). Using immunohistochemistry with antibodies against pre- and postsynaptic markers of excitatory synapses (vesicular glutamate transporter, vGLUT2; PSD-95), and super-resolution airyscan confocal imaging, we found tight colocalization between tdTomato+ axonal varicosities, vGLUT2 and PSD-95, an example of which is shown in **Fig. 4C**. This finding suggests that axons of thalamic neurons receiving cerebellar input form morphological synapses in the BLA. Axonal projections of DCN input-receiving thalamic neurons were also observed in other limbic regions including the nucleus accumbens core and shell (**Fig. 4D1,D2**) and anterior cingulate/prelimbic cortex (**Fig. 4D3,D4**).

**Figure 4.**
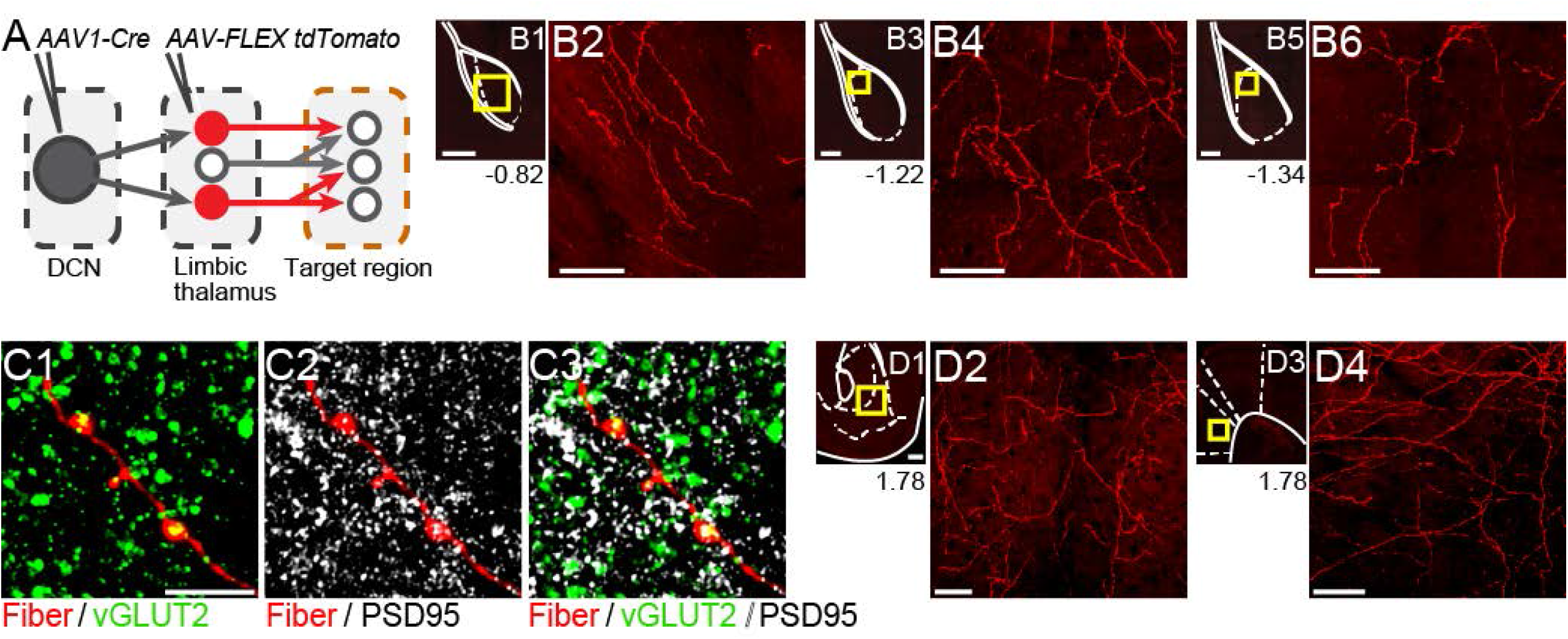
Thalamic neurons receiving cerebellar input form synapses in basolateral amygdala and also target the nucleus accumbens and prelimbic cortex. **A,** Schematic diagram of experimental approach. Targets of tdTomato+ axons of thalamic neurons receiving cerebellar input were identified through imaging. **B,** Mosaic confocal images of tdTomato+ axons along the anterior-posterior axis of the BLA. **C,** High resolution airyscan confocal images of tdTomato+ axons in the BLA colocalizing with presynaptic (vGLUT2) (**C1**) and postsynaptic (PSD95) (**C2**) markers of excitatory synapses. Green: vGLUT2, gray: PSD95, yellow/white in **C3**: overlay. **D,** tdTomato+ axons in nucleus accumbens (**D1,D2**) and prelimbic cortex (**D3,D4**). Yellow squares in B1,B3,B5 and D1,D3 show zoom-in areas for B2,B4,B6 and D2,D4 images, respectively. Numbers at bottom of images indicate distance (in mm) from bregma. Scale bars: B1,B3,B5,D1,D3: 200 μm; B2,B4,B6,D2,D4: 50 μm; C1-C3: 5 μm.

### 3.4 The centromedial and parafascicular nuclei emerge as functional nodes in cerebello-amygdala circuit

Our tracer overlap studies pointed to multiple thalamic nuclei as potential relays of cerebellar signals to BLA (Fig. 1E). Among them, the MD, CM and PF nuclei showed higher relative distribution of both BLA-projecting neurons and neurons that receive DCN input (Figs. 1D, 2C). For the remainder of this study, we focused on CM and PF nuclei and sought to substantiate their role as anatomical and functional relays of DCN-BLA connectivity through super-resolution microscopy and optophysiology.

Airyscan confocal imaging of slices from dual-tracer experiments (Fig. 1) revealed fluorescently labeled DCN axons (red) in contact with neurons that were retrogradely labeled from the BLA (green) in both CM (**Fig. 5A1,A2**) and PF (**Fig. 5A3–5**) nuclei. The existence of functional monosynaptic DCN-CM/PF connections was tested in the subset of electrophysiological experiments from Fig. 3 that targeted CM/PF neurons (**Fig. 5B**). Under basal conditions, CM/PF neurons received synaptic inputs from the DCN (at Vm = −70 mV; average amplitude ± SEM: −197.5 pA ± - 80.14, n = 6) (**Fig. 5C1,C5**) with short onset latency (average latency ± SEM: 2.4 ms ± 0.18) (**Fig. 5C6**), which is consistent with direct monosynaptic connections. Application of the sodium channel blocker tetrodotoxin (TTX) abolished the inputs (average amplitude ± SEM: −5.1 pA ± −2.03) (**Fig. 5C2,C4–5**), which recovered upon addition of the potassium channel blocker 4-AP (average amplitude ± SEM: −151.8 pA ± −39.52) (**Fig., 5C3–5**) (Friedman’s non-parametric repeated measures ANOVA: x^2^_r_ = 9, n = 6, p = 0.008; Dunn’s multiple comparison test: Baseline vs TTX: p = 0.02, Baseline vs TTX+4AP: p = 0.99), confirming their monosynaptic nature.

**Figure 5.**
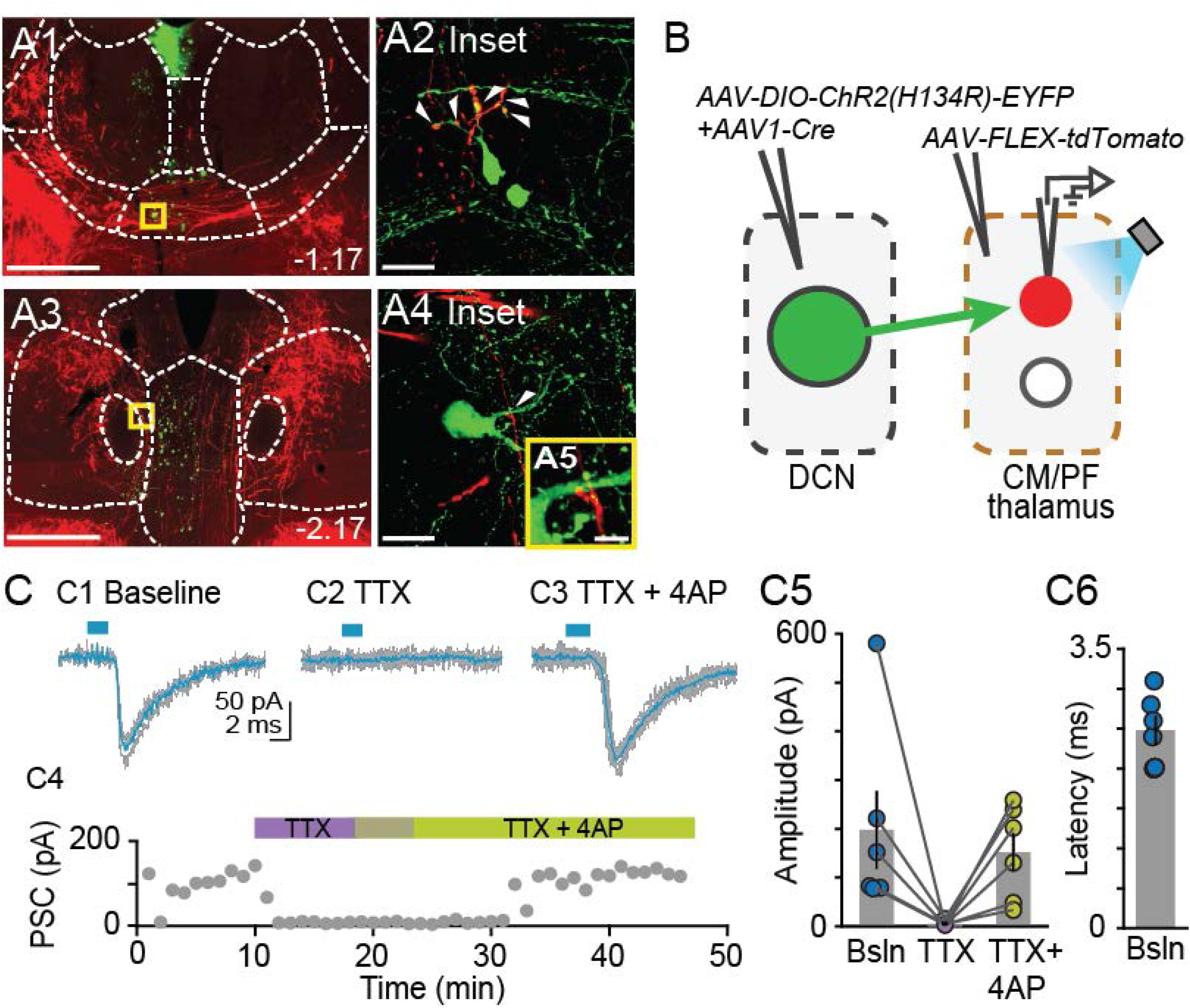
Centromedial and parafascicular neurons project to basolateral amygdala and receive functional monosynaptic input from the cerebellum. **A1-A4,** Airyscan confocal images of DCN axons (red) and BLA-projecting neurons (green) in the centromedial (CM; A**1**) and parafascicular (PF; A**3**) thalamic nuclei. **A2, A4-5,** zoomed-in areas in yellow squares from A1 and A3. Scale bars: A1,A3: 500 μm; A2,A4: 20 μm; A5: 5 μm. **B,** Schematic diagram of ex vivo optophysiology approach to test for monosynaptic connections between DCN and CM/PF thalamic n. **C1-C3,** Average ChR2-evoked synaptic current (teal), overlaid onto single trial responses (gray), at baseline (**C1**); upon addition of the action potential blocker tetrodotoxin (TTX, 1 uM) (**C2**); after further addition of the potassium channel blocker 4-aminopyridine (4AP, 100 uM) (**C3**). **C4**, Time course of wash-in experiment for the same example cell. **C5,** Summary of effects on amplitude (mean ± SEM) of ChR2-evoked synaptic responses for the indicated conditions. Bsln: baseline. **C6,** Average (± SEM) onset latency of ChR2-evoked responses at baseline.

Finally, we tested whether BLA is a target of DCN input-receiving CM/PF neurons (**Fig. 6**). We virally expressed ChR2 in DCN and stimulated cerebellar axonal projections in thalamic slices while recording from BLA-projecting CM/PF neurons (whole-cell voltage clamp mode, Vm = −70 mV), which were retrogradely labeled with CtB-CF568 in BLA (**Fig. 6A**). Optogenetic stimulation elicited reliable DCN-CM/PF synaptic responses (average amplitude ± SEM: −104.1 pA ± −37.1, n = 8) (**Fig. 6C,D1**) with short latency (3.35 ms ± 0.25) (**Fig. 6D2**). Combined with the imaging findings (Fig. 5), our electrophysiological results argue strongly for a DCN-BLA disynaptic circuit that recruits CM/PF nuclei as node.

**Figure 6.**
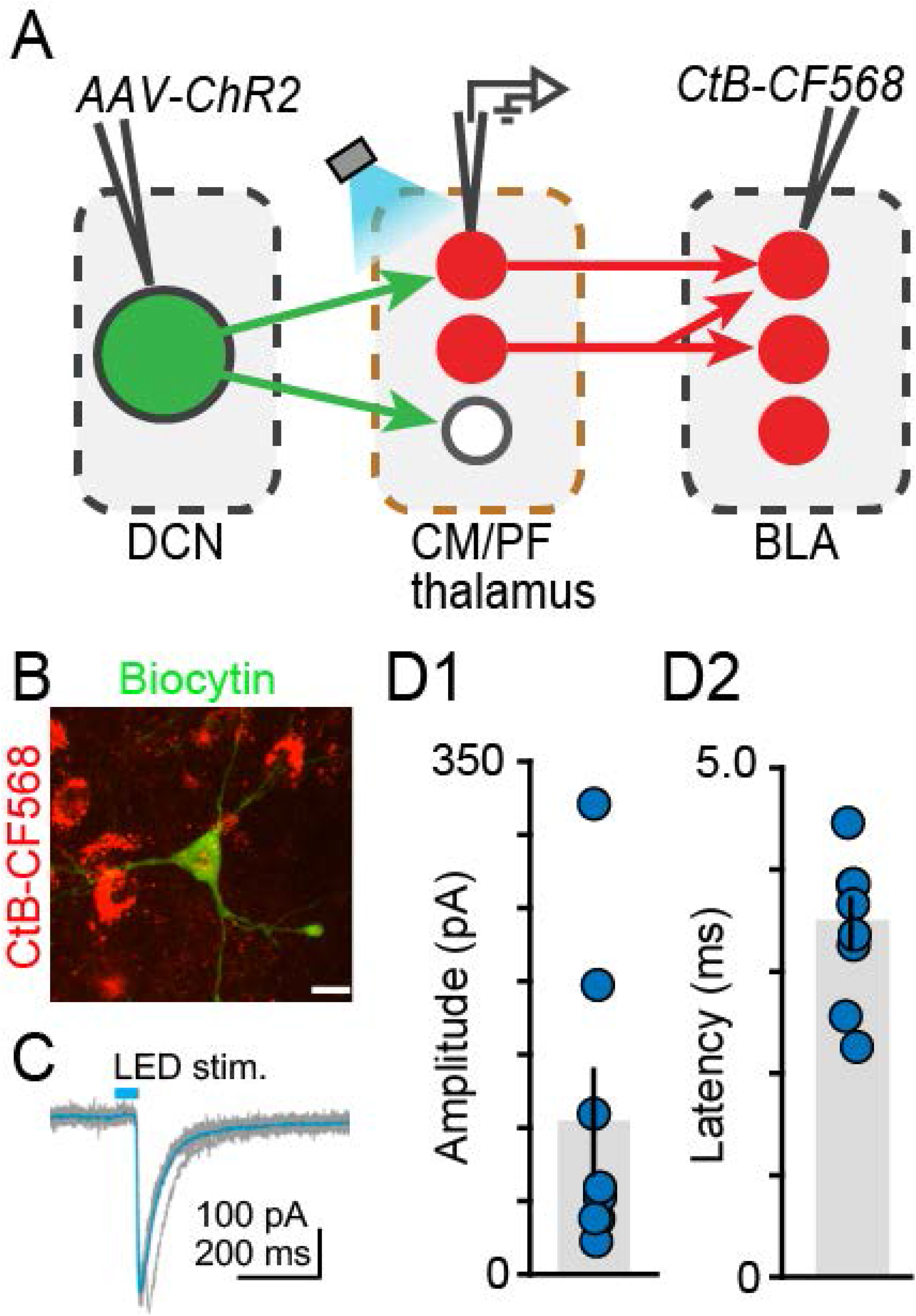
The centromedial and parafascicular thalamus is a functional node of the cerebello-amygdala circuit. **A,** Experimental approach. **B,** Example BLA-projecting neuron in centromedial (CM) thalamus retrogradely labeled with CtB CF-568 (red) is also labeled with biocytin (green) through the patch pipette. Scale bar = 10 μm. **C,** Example ChR2-evoked synaptic response. Average trace (teal) overlaid onto single trials (gray). **D1,D2,** Average (± SEM) amplitude (D1) and onset latency (D2) of ChR2-evoked synaptic currents at DCN-CM/PF synapses.

## 4 Discussion

Cerebellar connections with the amygdala have been posited previously but the neuroanatomical substrate of this connectivity has been elusive ^1,5,53^. Here, we obtained insight into cerebello-amygdala circuitry by combining various tracing approaches with advanced imaging and optophysiology. We established the existence of a disynaptic circuit between cerebellar nuclei and BLA, thus providing the first blueprint of cerebello-amygdala connectivity at the mesoscale level. The circuit recruits at least the centromedial and parafascicular thalamic nuclei (Figs. 5,6), and most likely also other nuclei of the limbic thalamus (Fig. 1), as relay nodes. In addition, we identified the intralaminar thalamic cluster and MD nucleus as recipients of the majority of cerebellar inputs to limbic thalamus (Fig. 2). Finally, and in addition to BLA, we identified axonal projections of DCN input-receiving thalamic neurons in limbic regions such as nucleus accumbens core and shell and anterior cingulate/prelimbic cortex (Fig. 4).

### 4.1 The limbic thalamus as a target of cerebellar inputs

We targeted the limbic thalamus as a conduit of cerebello-amygdala communication because several of its nuclei foster BLA-projecting neurons in close proximity to DCN axons (Fig. 1). DCN projections to limbic thalamus have been observed before ^54–58^ but the existence of functional synaptic terminals has only been validated for centrolateral and PF intralaminar nuclei ^30,52^, and never on amygdala-projecting neurons. Our optophysiological experiments also provided first evidence for the presence of active synaptic connections (as opposed to just passing axons) in paracentral and centromedial (part of intralaminar group), intermediodorsal and rhomboid (part of midline group), and mediodorsal nuclei (Fig. 3), expanding the repertoire of non-motor cerebellar targets and paving the way for causal manipulations.

### 4.2 Technical considerations

To chart cerebello-amygdala neuroanatomical connections, we used powerful circuit mapping tools including anterograde and retrograde tracer viruses and the transneuronal AAV1-Cre approach ^50,59–61^. A distinct advantage of our approach, which combined AAV1-Cre with viral injections of conditionally expressed fluorescent tracers (as opposed to reporter mouse lines), is the ability to definitively point to the thalamus as the source of the axonal projections in BLA, NAc and prelimbic cortex- as opposed to e.g., the VTA, which also receives DCN inputs and projects to these regions ^62–66^. Thus, our approach enabled conclusive interpretation of anatomical connectivity results. On the flip side, injection coverage/spill and viral tropism ^61^ need to be considered. Tropism, in particular, could skew interpretation of disynaptic inputs, as some cell groups in the limbic thalamus might be more efficiently infected by AAVs. Tropism could also explain why recent efforts to trace di- and tri-synaptic cerebellar efferent pathways with herpes simplex viruses did not identify the CM/PF pathway to BLA ^67^. Lastly, one potential concern could be the propensity of AAVs to be transported in the retrograde direction at high titers ^50,68^. To remediate these concerns, we used strict inclusion criteria for injection sites; employed a combination of viral and non-viral anterograde and retrograde tracers; optimized viral titers so as to minimize retrograde transport; and confirmed circuit connections with slice optophysiology.

### 4.3 Proposed functions of the DCN-BLA circuit

Our discovery of the DCN-BLA connection through CM/PF provides an essential map for future investigation of circuit function. The circuit, which could account for the previously observed short-latency cerebello-amygdala responses ^37^, could convey cerebellar information about prediction, salience and/or valence to BLA, shaped by the intrinsic, synaptic and integrative properties of the nodes. Indeed, the cerebellum is known to encode such information ^8,69–73^, which is also seen in BLA ^35,74–79^, and which is thought to be used by CM and PF during aversive conditioning, observational learning and reward-seeking behavior ^30,46,80–83^.

The cellular targets of cerebello-thalamic axons in BLA remain to be determined but likely include at least BLA principle neurons, which are the major recipients of CM input ^84^. The patterns of BLA ensemble activity triggered by distinct cerebello-thalamic inputs could serve different aspects of cerebellum-relevant emotional functionality, which includes modulation of anxiety and learned fear ^29,32,85–88^; processing of facial emotional expressions ^89,90^; regulation of emotional reactivity ^91,92^; and even reward-driven motivated behavior ^27,31,93,94^.

The BLA is not the sole nucleus in the amygdala complex that receives cerebellar signals ^95^. Similarly, it is unlikely that the CM and PF are the only nuclei to serve cerebello-amygdala communication (our findings; and ^96^). Further studies are warranted to delineate the complete neuroanatomical and functional landscape of cerebello-amygdala connectivity. Our findings constitute the first step toward this goal.

## 6 Conflict of Interest

The authors declare that the research was conducted in the absence of any commercial or financial relationships that could be construed as a potential conflict of interest.

## 7 Author Contributions

SJJ, KV and DF designed the study; SJJ, KV, AD, AP and YI performed experiments; SJJ, KV, EA and DF analyzed data; MB, EPF, JV, MFF, and MA assisted with cell counting; SJJ, KV, AD and DF wrote the manuscript with input from authors.

## 8 Funding

This work was supported by R21MH114178, NSF1754831, a NARSAD Young Investigator Grant, Brain Research Foundation grant BRFSG-2017-02, and a Whitehall Foundation research award to DF; a NARSAD 2018 Young Investigator Grant to EA; a UC Davis Provost’s undergraduate fellowship to MA; a NIMH T32MH112507 fellowship to KV; and a NIH T32GM007377 and a UC Davis Dean’s Distinguished Graduate Fellowships to AD.

## 9 Acknowledgments

We thank Dr. Brian Wiltgen of UC Davis for access to imaging equipment; and Fioravante lab members for comments on a previous version of the manuscript.

## Notes

### Competing Interest Statement

The authors have declared no competing interest.

